# Functional Data Clustering of Filopodia Dynamics

**DOI:** 10.64898/2026.11.04.764394

**Authors:** Nneka Enumah, S.H Dinuka Sewwandi De Silva, Hannah Cho, Gabriella Schuckers, Ginger Hunter, Emmanuel Asante-Asamani

**Affiliations:** Department of Mathematics, Clarkson University, 8 Clarkson Avenue, Potsdam, 13699, NY, USA; Department of Biology, Clarkson University, 8 Clarkson Avenue, Potsdam, 13699, NY, USA; Department of Biology, Howard University, 2400 6th St NW, Washington, 20059, DC, USA; Department of Mathematics, Saint Lawrence University, 23 Romoda Dr, Canton, 13617, NY, USA; Department of Mathematics, Alma College, 614 W Superior St, Alma, 48801, MI, USA

**Keywords:** Functional data clustering, filopodia dynamics, bristle patterning, drosophila melanogaster

## Abstract

Notch signaling plays a significant role in tissue patterning. During patterning of the fruit fly dorsal thorax peripheral nervous system, basal filopodia extend the spatial range of cell-cell communication, yet it is still unclear how Notch is activated in long range signaling via filopodia interaction. Filopodia undergo dynamic length changes during their lifetime, which could supply the mechanical pulling force required for notch activation. However, their dynamics are often characterized using individual scalar values such as maximum length and extension rate which could obscure more intricate dynamics. Our hypothesis is that signaling filopodia exhibit distinct and recurrent temporal patterns which can be identified by a consideration of their complete lifetime trajectories. Our work analyzes 60 filopodia trajectories at the basal surface of the fly dorsal thorax during patterning. Length profiles were smoothed using cubic B-splines and clustered by first reducing them to their dominant functional principal component followed by the application of various clustering algorithms. We found that 94.29% of the overall trajectory variation is explained by the first four functional principal components which account for variations in maximal length, extension and retraction rate, duration of stationary phase and length fluctuation frequency. Five recurrent temporal profiles (dynamic extension modes) with varying trajectory shapes and physical attributes, were identified using trimmed K-means. These findings demonstrate that whereas the major source of structural variation in filopodia profiles remains the length, extension rate and retraction rate, nontrivial combinations of these create distinct dynamic modes of extension which provide new insight into their length profiles. Our work thus offers a quantitative framework for future research connecting filopodia dynamic extension modes to productive Notch activation.

## 1. Introduction

Notch-signaling is a highly conserved juxtacrine signaling pathway in which a physical interaction between the Notch receptor in a signal receiving cell and Deltaligand in a signal sending cell results in the cleavage of the Notch receptor and translocation of the Notch intracellular domain (NICD) into the nucleus of the signal receiving cell [1]. These signaling events are well known to regulate lateral inhibition, which is a key mechanism for coordinating developmental processes in most organs and tissues in metazoans [2, 1, 3, 4]. Generally, a high level of Delta in one cell results in the suppression of delta expression in adjacent cells[5]. A dysregulation of this signaling pathway has been associated with the onset and progression of hematologic malignancies and solid tumors, as well as genetic disorders such as Adams-Oliver syndrome and Alagille syndrome [6, 7]. Drosophilia melanogaster is a genetically tractable model organism for studying Notch-Delta signaling (ND signaling). Of particular interest is the use of ND signaling in the organization of sensory bristles on its dorsal thorax during pupal development. In this process, initially bipotential cells with relatively uniform levels of Notch and Delta progressively increase or decrease Notch levels via lateral inhibition. Eventually, cells that achieve higher levels of Delta become sensory organ precursor (SOP) which later develop into bristles, while those with high levels of Notch become epithelial cells. Recent experimental and computational simulations have suggested that long-range ND signaling, mediated by signaling filopodia (cytonemes), is a necessary addition to traditional junctional signaling in order to explain the observed spacing between SOP cells[8]. Although Notch receptors and Delta ligands have been observed to localize along filopodia [9], the mechanism by which Notch receptors are activated along filopodia is unknown.

Notch activation requires mechanical pulling force which stretches the receptor and reveals a cleavage site where enzymatic activity separates the extracellular from intracellular domain (NICD). This pulling force is provided by endocytosis of Delta ligands into signal sending cells, when cells are in adjacent contact [10, 11, 12]. While endocytosis has been recently observed to occur along filopodia through clathrin coated pits [13, 14], it is unknown whether this mechanism is at play for filopodia in the dorsal thorax [10]. Alternatively, filopodia in the dorsal thorax have been observed to be highly dynamic, undergoing cycles of protrusion and retraction with an average lifetime of about 10 minutes [15]. Additionally, the filopodia network has been observed to be under mechanical tension, undergoing a rapid recoil when the network is cut in local regions [15]. Based on this, it stands to reason that the pulling force from the protrusion-retraction dynamics of filopodia can supply sufficient mechanical force to activate Notch. While distributions of filopodia characteristics such as maximal length, lifetime, retraction rate and extension rate have been quantified in the Drosophila notum[15] and other model systems [16, 17, 18, 19], we have very little understanding about cytoneme length versus time dynamics within a single lifetime. Such knowledge will be critical to quantitatively assessing the likelihood of Notch activation within the lifetime of filopodia.

Measurements of filopodia characteristics such as maximal length, lifetime, extension and retraction rates, while informative, do not reliably indicate signaling activity. For instance, increased filopodia length has been shown to co-exist with a reduced Notch-transcriptional response [20]. More broadly, these scalar characteristics obscure more intricate behaviors within the lifetime of the filopodia. As a result, two filopodia with significantly distinct temporal trajectories can have comparable scalar descriptions. Recent work by González-Mendez et al. differentiated between cytonemes that extended and retracted immediately and those that stayed extended for a while before retracting [19]. Other studies have attempted to make similar characterizations [21, 22]. Beyond these, it is unclear whether filopodia can undergo other intricate length dynamics and what unbiased framework can be used to determine these dynamic modes. We hypothesize that signaling filopodia exhibit distinct recurrent patterns that can be characterized through an observation of their full temporal trajectory. We test this hypothesis within the theoretical framework of functional data analysis, where experimental temporal length measurements are assumed to be discrete samples from an underlying continuous function [23, 24, 25]. Since our interest is in classifying dynamic length modes within a single filopodia lifetime, we employ functional data clustering to group smooth functions fit to length versus time profiles normalized with respect to lifetime. Functional groups are identified by quantifying the primary variation within a collection of functions using functional principal component analysis. These methods have found applications in the classification of gene-expression curves and other longitudinal biological data [26, 27, 28], but have not been applied to characterize dynamic filopodia modes. In this work, we will use functional data clustering to perform an unbiased trajectory shape analysis of filopodia data obtained from the dorsal thorax of developing fruit flies, without first reducing each protrusion to a limited set of scalar summary statistics. Identification of clear dynamic filopodia modes from our analysis will facilitate future model driven analysis of the likelihood of Notch activation within the lifetime of a filopodium.

The rest of the paper is structured as follows. The preprocessing, functional representation, functional principal component analysis, clustering techniques, and characterization of the final filopodial clusters are all covered in the Materials and Methods section. The Results section illustrates the key types of trajectory variation, compares the clustering methodologies, and analyzes the dynamic and physical characteristics of the identified clusters. The key findings and their implications are summarized in the Discussion and Conclusion.

## 2. Materials and Methods

Live confocal microscopy of Drosophila filopodia. Fly strains used in this study: neuralized-GAL4, UAS-GFP-moesinCA / TM6BTb (Hunter Lab) and UAS-LexA-RNAi (Bloomington Drosophila Stock Center 67947). neuarlized-GAL4 expresses GAL4 in neural precursor (SOPs) only. UAS-GFP-moesinCA is the constitutively active F-actin binding domain of Moesin tagged with GFP under control of the UAS promoter. These lines were crossed to generate F1 pupae for imaging. F1 0 hAP (hours after pupariation) white prepupae were selected then aged to 12hAP. Pupae were then mounted on a glass slide with double sided tape and the pupal case is removed [29]. The prepared samples are imaged on an Leica DMi8 inverted line scanning confocal microscope using an 40x/0.8NA objective and Leica LASX software. All images were subsequently processed and analyzed using FIJI/ImageJ.

### 2.1. Preprocessing

#### Time Normalization

Let *y_ij_* denote the observed length of filopodium *i* at time *t_ij_*, where *i* = 1*, . . . ,* 60 and *j* = 1*, . . . , n_i_* is the observation within that filopodium’s trajectory. To compare trajectories with different lifetimes, the observed time in minutes was mapped to a common interval [0, 1]:

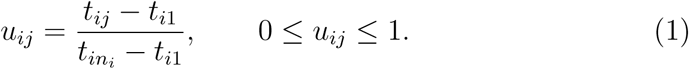

Thus for each filopodium, *u* = 0 represents the first observation and *u* = 1 represents the last observation f. This representation preserved the original length scale while eliminating variations in life time duration from the functional form analysis. For the computation of lifetime distribution,extension and retraction rates, and oscillation frequencies the observed time was maintained independently.

#### Data smoothing with B-splines

Each trajectory was represented as a linear combination of a common set of cubic B-spline basis functions:

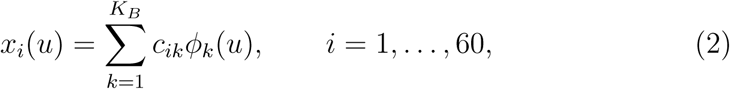

where *ϕ_k_*(*u*) is the *k*th shared basis function and *c_ik_* is its coefficient for trajectory *i*. Seven basis functions were used (*K_B_* = 7), matching the number of observations in the shortest trajectory and ensuring that the same finite-dimensional representation could be estimated for every filopodium. The coefficient vector

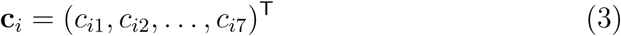

therefore provided a seven-parameter representation of the complete trajectory. Coefficients were estimated by fitting the shared basis to the observed lengths on the normalized time grid:

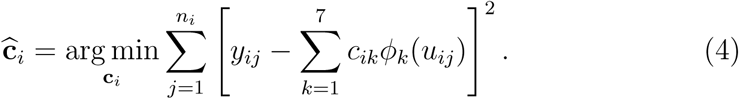

The fitted functions were evaluated visually against the observed data to verify that the seven-function basis maintained the primary extension, maximum length,retraction and local-undulation properties without duplicating point-to-point measurement noise.

### 2.2. Functional PCA analysis (FPCA)

FPCA was applied to the fitted functions to identify orthogonal modes of variation among filopodia length trajectories. The sample mean function is taken as

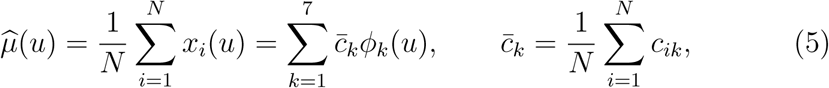

with *N* = 60. The empirical covariance surface was defined as

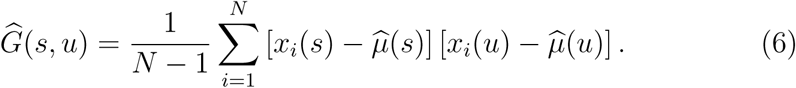

Eigenfunctions *b_v_*(*u*) and eigenvalues *λ_v_*were obtained from

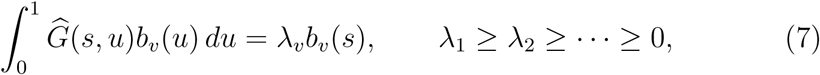

subject to the orthornomality constraint

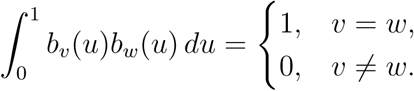

The score of trajectory *i* on component *v* was

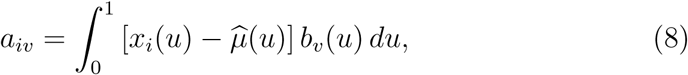

and the truncated reconstruction was

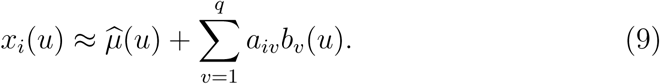

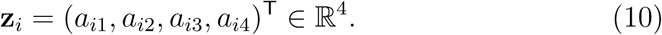

Plotting the mean curve along with the perturbations 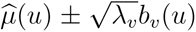 allowed us to determine the significance of each component. This allowed for the interpretation of differences in overall length, extension and retraction timing, stabilisation behaviour, and localised fluctuations in relation to separation in score space.

### 2.3. Functional Data Clustering Techniques

The four FPCA scores representing each filopodium were clustered, and the dissimilarity between the score vectors - two filopodia *i, j* - was measured using the Euclidean distance,*d*(*i, j*) where smaller distances indicates similar trajectory pattern and vice-versa.

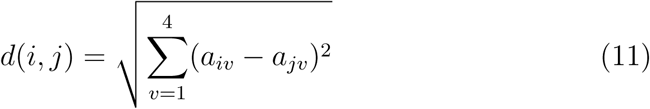

The within-cluster sum of squares, mean silhouette coefficient, and biological interpretability of the resulting mean trajectory profiles were used to assess a range of cluster sizes, from two to ten.

#### 2.3.1. K-means

The FPCA score vectors were divided into *K* disjoint clusters *C*_1_*, . . . , C_K_* by K-means clustering by minimizing

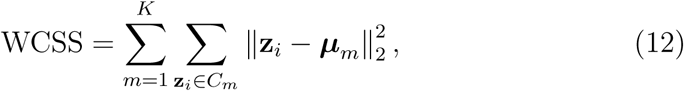

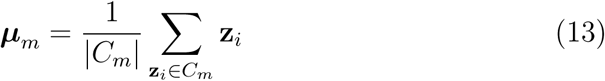

where *µ_m_*is the centroid of cluster *m*. After initialization, centroids were recalculated from the allocated observations, observations were assigned to their closest centroid, and this process was repeated until there were no further changes in cluster membership.

#### 2.3.2. Trimmed K-means

In order to prevent a small number of unusual trajectories from unduly impacting the centroids, trimmed K-means was employed as a reliable substitute. A trimming fraction of *α* = 0.05 was applied. The three trajectories with the biggest nearest-centroid distances were therefore left unassigned, and *h* = *N* (1 − *α*) = 57 trajectories were kept in the fitted partition. If the kept index set is denoted by *H* ⊂ {1*, . . . , N* }, the trimmed objective was

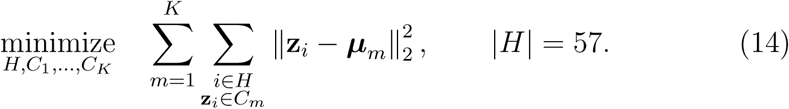

Every trajectory was temporarily allocated to its closest centroid at each iteration, the centroids were recalculated using only the retained observations, the nearest-centroid distances were ordered, and the largest 5% were cut. The process was repeated until the cluster assignments and the retained set stabilised.

#### 2.3.3. Agglomerative hierarchical clustering

Each FPCA score vector was initially used as a singleton cluster in agglomerative hierarchical clustering, which then combined the closest two clusters until only one cluster remained. To get partitions that were directly comparable to the centroid-based solutions, dendrograms were chopped at *K* = 5. Four definitions of linkage were considered. For clusters *A* and *B*, single linkage distance is defined as:

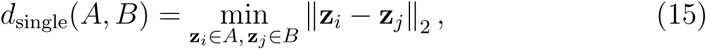

complete linkage:

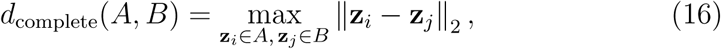

average linkage:

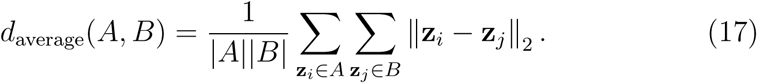

Ward linkage merged the pair producing the smallest increase in within-cluster dispersion

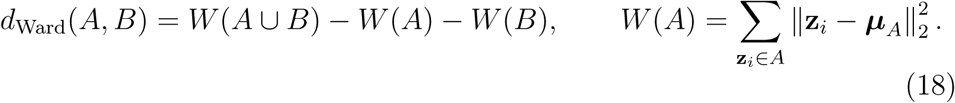

Because linkage heights have method-specific scales, merge distances were interpreted within each dendrogram and were not compared numerically across linkage definitions.

### 2.4. Characterization of Filopodia Clusters

Both the Calinski-Harabasz Index and the silhouette score were used to assess cluster quality. Let *a*(*i*) be the mean distance of observation *i* to the other members of its assigned cluster, and let *b*(*i*) be the minimum mean distance of *i* to any other cluster. The silhoutte score is given by

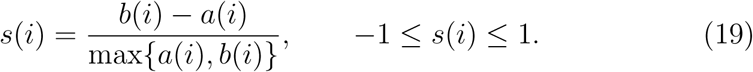

The Calinski-Harabasz index was calculated as

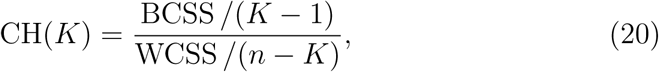

where

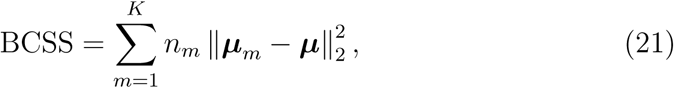

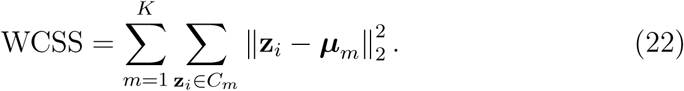

Here, *n_m_* is the size of cluster *m*, ***µ****_m_* is its centroid, and ***µ*** is the overall centroid. Higher silhouette and Calinski-Harabasz values indicate more compact and better-separated clusters.Both indices were assessed on the 57 trajectories that were kept for trimmed K-means in order to prevent unassigned data from skewing the internal validity metrics.

#### 2.4.1. Estimation of Extension/Retraction Rate

A time-weighted average of the rates computed between successive observations was used to estimate the mean extension and retraction rates of each filopodium. Let *y_ij_* denote the observed length of filopodium *i* at time *t_ij_*, where *j* = 1*, . . . , n_i_*. Length measurements were recorded at intervals of 30 seconds (Δ*t* = 0.5 min).

The length change and the associated time interval for successive measurements were defined as

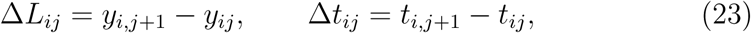

where *j* = 1*, . . . , n_i_ −* 1.

The instantaneous interval-based rate was estimated as

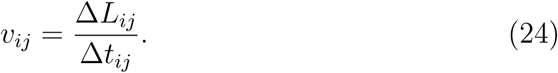

Positive values of *v_ij_* indicate extension, whereas negative values indicate retraction. Consequently,

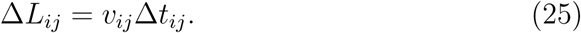

To calculate the mean extension rate, only intervals with positive length changes (Δ*L_ij_ >* 0) were considered. Let

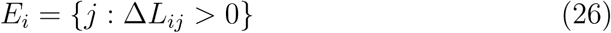

denote the set of extension intervals for filopodium *i*. The mean extension rate was calculated as the time-weighted average

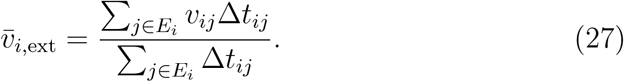

Equivalently,

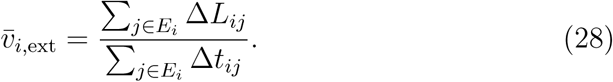

Similarly, the mean retraction rate was calculated using only intervals in which the filopodium decreased in length. Let

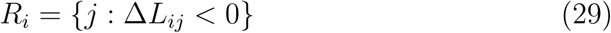

denote the set of retraction intervals. The mean retraction rate was defined as

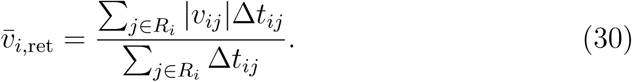

Equivalently,

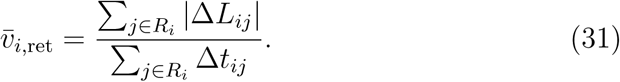

Retraction rates were expressed as positive magnitudes.Both computations did not include intervals with no change in length (Δ*L_ij_* = 0). A filopodium’s associated rate was deemed undefined if it lacked extension or retraction intervals. Using the initial observation times, the mean extension and retraction rates were independently calculated for every filopodium. The mean and standard deviation were then used to summarize the resulting rates within each cluster.

#### 2.4.2. Maximal Length Distribution and Average Lifetime Distribution

Maximum length and lifetime were calculated separately for each filopodium using the original observations on the observed-time scale. For filopodium *i*, the maximum observed length was defined as

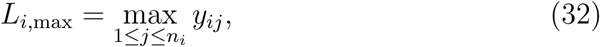

where *y_ij_* is the observed length at measurement *j* and *n_i_* is the total number of measurements for filopodium *i*. The lifetime of filopodium *i* was calculated as the elapsed time between its first and final observations:

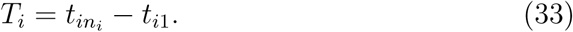

Because observations were recorded at 30-second intervals, lifetime could equivalently be calculated in minutes as

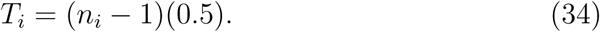

Thus, lifetime represented the period during which the filopodium was continuously observed, from its first detected appearance to its final recorded observation.

The distributions of maximum length and lifetime were examined within each cluster. For cluster *m*, containing *n_m_* filopodia, the mean maximum length was calculated as

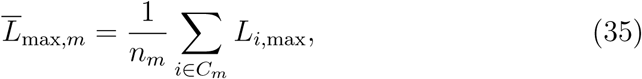

and the corresponding standard deviation was

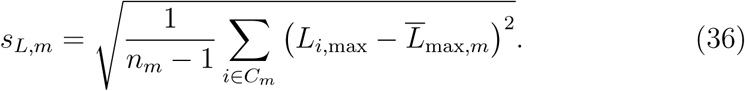

Similarly, the mean lifetime within cluster *m* was

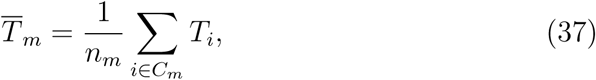

with standard deviation

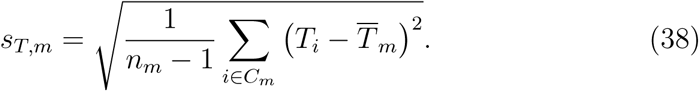

To describe variations in filopodial size and persistence, the cluster-specific distributions of maximum length and observed longevity were analysed. Numerical summaries were supplied via cluster-level means and standard deviations, while boxplots displayed the distribution of individual values.

#### 2.4.3. Power Spectral Analysis

Power spectral analysis was performed separately for each filopodium to characterize the frequency content of its length fluctuations. The observation times were normalized to the interval [0, 1], and the sampling interval, Δ*u_i_*, was calculated as the median difference between consecutive normalized observation times. A least-squares B-spline was fitted to the valid length measurements and evaluated on a uniform grid from zero in increments of Δ*u_i_* up to the largest grid point not exceeding one. The number of basis functions was the smaller of the requested basis count and the number of valid observations. The resulting fitted sequence was used for spectral analysis, with a normalized sampling frequency of *f_s,i_* = 1*/*Δ*u_i_*.

For filopodium *i*, let *x_ij_* denote the fitted length at evaluation point *j*, and let *N_i_* denote the number of evaluation points. The sequence was centered by subtracting its mean:

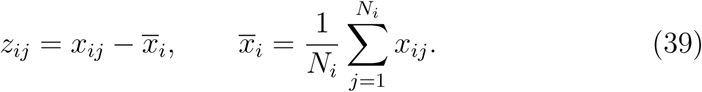

Centering removed the constant contribution of mean length so that the spectrum described variation around that mean. The discrete Fourier transform was computed using the fast Fourier transform:

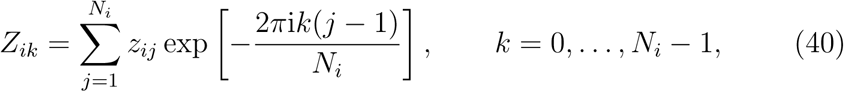

where i denotes the imaginary unit. The corresponding nonnegative frequencies were

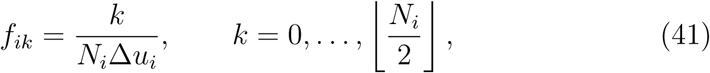

expressed in cycles per observed lifetime. The one-sided power spectrum was calculated as

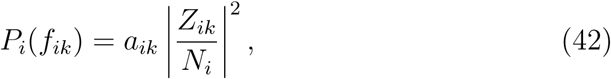

where, in the implementation used, *a_ik_* = 2 for interior retained frequency bins and *a_ik_* = 1 for the zero-frequency and final retained bins. The resulting values represented squared-amplitude power per bin rather than power spectral density. The normalized Nyquist frequency was *f_s,i_/*2, and the frequency-bin spacing was 1*/*(*N_i_*Δ*u_i_*). The dominant frequency was defined as the positive frequency with the greatest spectral power:

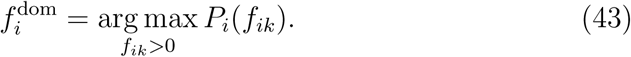

Its corresponding period was 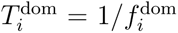, expressed as a fraction of the observed lifetime. When multiple bins shared the maximum power, the lowest tied frequency was selected. The power-weighted mean frequency was also calculated, giving greater weight to frequencies with higher spectral power:

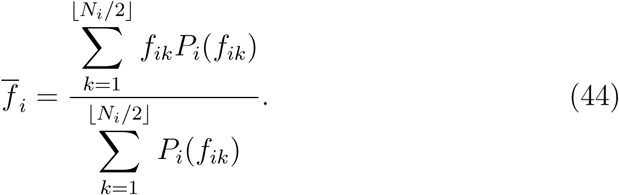

Dominant frequencies, corresponding periods, and power-weighted mean frequencies were calculated only for trajectories with positive nonzero-frequency power.

## 3. Results

We analyzed 60 distinct filopodial length trajectories. Over the course of each filopodium’s observable lifetime, each trajectory comprised a series of length measurements in micrometres *µm* which was taken at intervals of thirty seconds (see Figure 1A). The trajectories varied in lifetime and included seven to ninety observations. The initial and last observations were aligned without altering the measured length by mapping each trajectory to the relative-time interval [0, 1]. This step made it possible to compare trajectories with varying lifetimes based on the times that extension, maximum length, and retraction took place relative to their individual lifetimes (see Figure 1 B). See the methods section for details on the time normalization. From the initial recorded observation to the final recorded observation, the entire trajectory was considered a single functional observation. Each trajectory was smoothed using a shared cubic B-spline basis defined over the same relative-time domain. Figure 1 C shows the B-spline representation for the length profile in Figure 1B. The set of B-spline basis functions used for smoothing is shown in Figure 1D.

**Figure 1:**
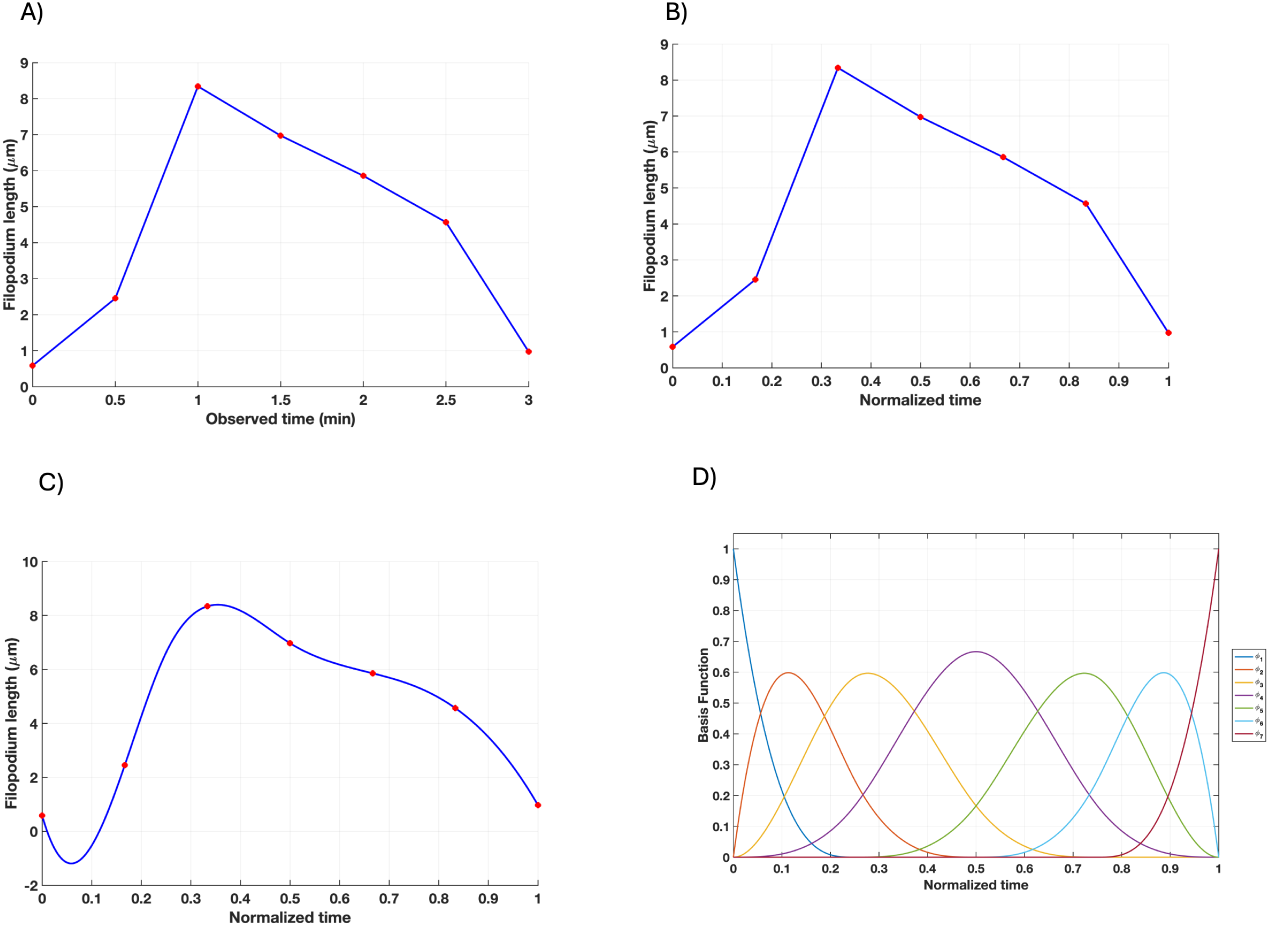
Preprocessing and functional representation of a representative filopodial length trajectory. (A) Measurements of length using the initial observation-time scale. (B) The same measurements were standardized to the interval [0, 1], where 0 and 1 represent the initial and final observations, respectively. (C) Cubic B-spline fit to the normalized measurements; the continuous fitted trajectory is shown by the blue curve, and the observed lengths are indicated by red markers. (D) All trajectories on the same normalized-time domain are represented by seven cubic B-spline basis functions. During time normalization, the length stayed in µm.

The prominent patterns of variation among the smoothed curves were subsequently found using functional principal component analysis (FPCA), which also produced a low-dimensional score representation. The scores where clustered using K-means, trimmed K-means, and four agglomerative hierarchical clustering techniques. Then the mean functional profiles, lifetime and maximal-length distributions, extension and retraction rates, and frequency-domain behaviour were used to describe the resulting clusters. For parameters that rely on elapsed time, such as lifetime, extension and retraction rates, and oscillatory frequency, physical time was maintained separately.

### 3.1. Principal Variation in Filopodia Profile

FPCA was used to reduce the correlated variation among the smoothed trajectories to a small set of orthogonal shape features. Details of this process are discussed in the methods section. The first four components explained 56.83%, 19.13%, 12.20%, and 6.14% of the total functional variation, respectively, giving a cumulative explained variance of 94.29%. PC1 was the dominant source of variation, while PCs 2–4 captured progressively smaller differences that were not represented by PC1. Retaining four components reduced each seven-coefficient spline representation to four uncorrelated scores while preserving substantially more variation than the first three components alone, which retained approximately 88.2%.

The mean function shows a broad extension phase followed by a maximum near the middle of normalized lifetime and retraction toward the endpoint (Fig. 2). The ±1-standard-deviation perturbations indicate that the components represented more than uniform differences in length. PC1, shown in Fig. 2A, primarily captures the overall amplitude or length variation i.e overall protrusion magnitude. PC2, shown in Fig. 2B, captures variations in the time at which maximum length is reached, which is driven by variations in the extension and retraction rates. PC3, shown in Fig. 2C, captures the variations in the duration of the stationary phase where filopodia length remains steady. PC4, shown in Fig. 2D, described more localized fluctuation, particularly around the early-to-middle and late portions.

**Figure 2:**
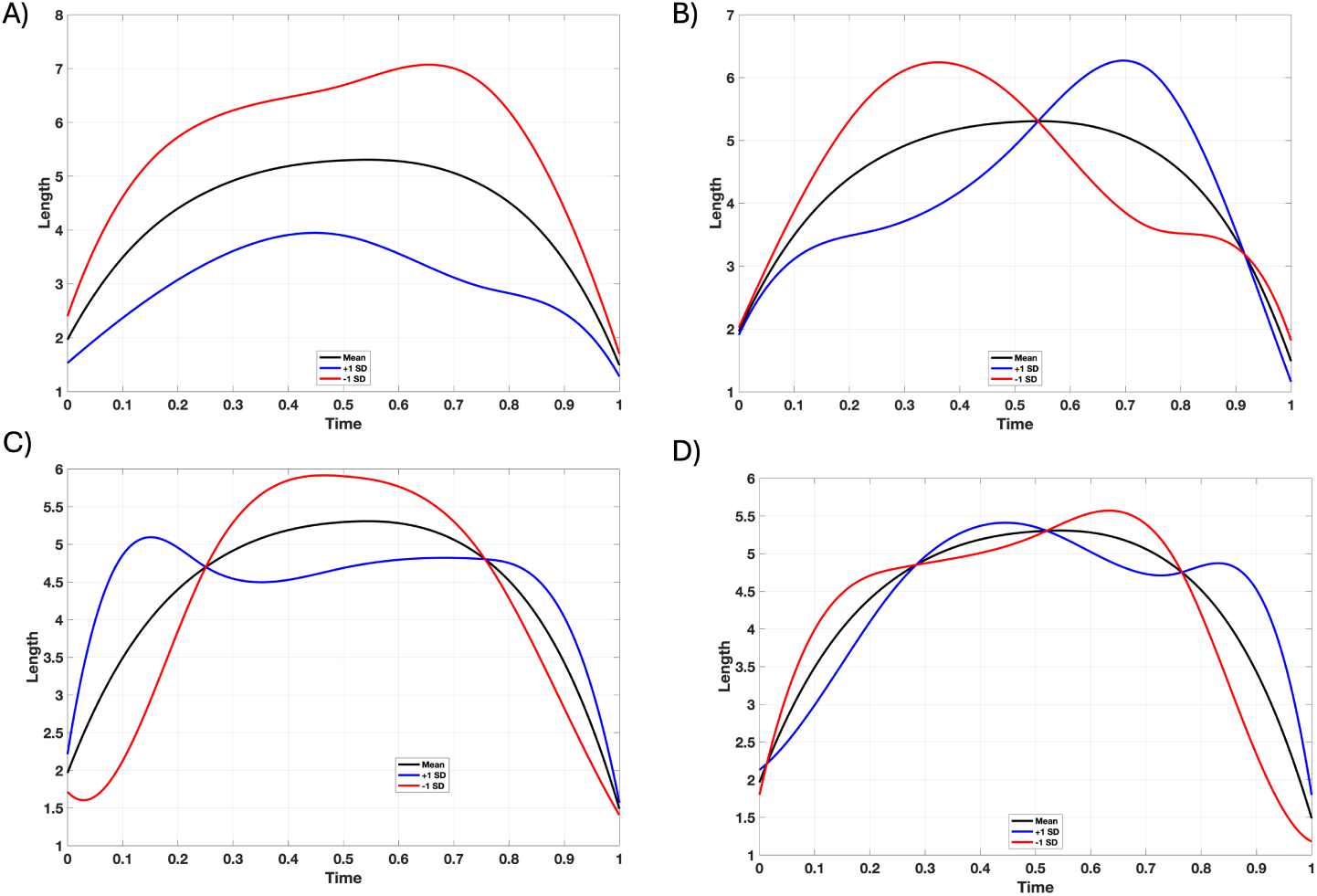
Functional principal component modes of filopodial length variation. The black curve in each panel is the sample mean function, and the blue and red curves show the mean perturbed by one standard deviation along the corresponding eigenfunction, 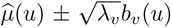. A)PC1 PC1–PC4 explained 56.83% of total variation. B)PC2 explained 19.13% of total variation C)PC3 explained 12.20% of total variation and D)PC4 accounted for 6.14% of total variation,(94.29% cumulatively). Separation between the perturbation curves identifies the normalized-time regions contributing most strongly to each component; crossings indicate changes in the direction of the component’s effect over the trajectory.

### 3.2. Trimmed K-means provides optimal clustering of filopodia

The four retained FPCA scores were clustered so that similarity was determined by complete trajectory shape rather than by any single measurement. K-means, trimmed K-means, and four hierarchical linkage rules were compared at *K* = 5 to determine whether the identified groups were robust to the clustering criterion. We choose *K* = 5 because the within cluster sum of squares (WCSS) curve, defined in see Eq. 12, showed an elbow, after which more clusters resulted in relatively minor decreases in within-cluster variation. Additionally, the five-cluster setup produced unique, physiologically understandable trajectory patterns while maintaining a good silhouette coefficient. The hierarchical solutions differed substantially across linkage definitions (Fig. 3). Single linkage produced pronounced chaining: most trajectories entered one large cluster while a few trajectories remained as small or singleton groups. Average linkage showed a similar, although less extreme, imbalance. Complete and Ward linkage produced more internally structured partitions, but cluster membership and the relative sizes of the five groups still varied with the linkage rule. Ward-linkage merge distances remained comparatively low through most agglomerative steps and then increased sharply during the final merges. This increase indicates that the last merges combined groups that were considerably more dissimilar than those joined earlier and supports retaining a multi-cluster representation.

**Figure 3:**
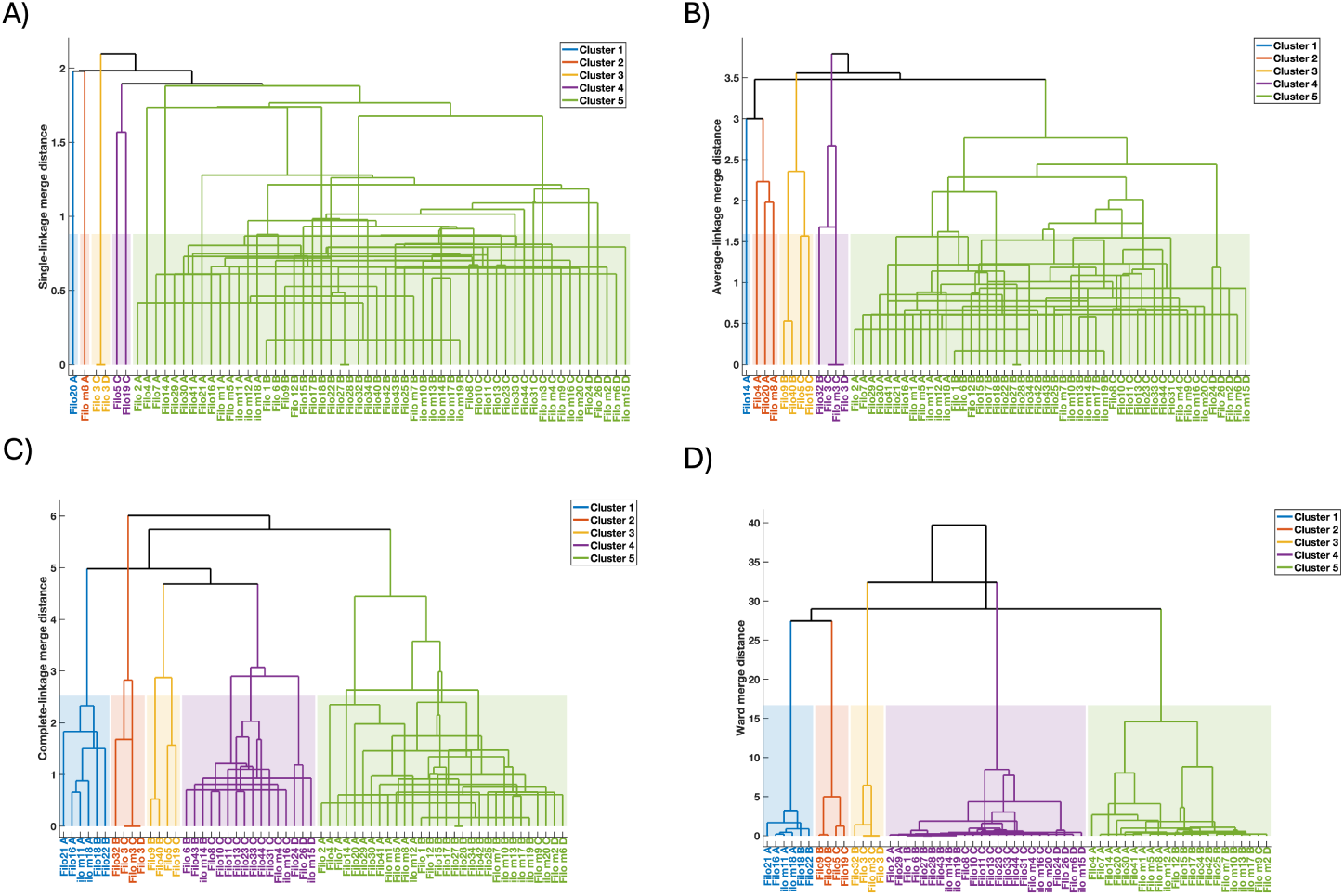
Agglomerative hierarchical clustering of the four-dimensional FPCA score vectors using four linkage definitions. Dendrograms show single, average, complete, and Ward linkage in reading order. Branch height gives the method-specific merge distance, and colors indicate the five-cluster partition obtained by cutting each dendrogram. Shaded blocks identify the resulting cluster memberships. Merge heights should be interpreted within, rather than numerically compared across, linkage methods. Single and average linkage place most trajectories in one large cluster and isolate several small groups, whereas complete and Ward linkage yield more structured partitions.

At *K* = 5, trimmed K-means had the largest mean Calinski-Harabasz index (0.3308), followed by K-means (0.3186) and Ward linkage (0.2742) (Table 1). Single linkage had the smallest value (0.0783), consistent with the pronounced chaining seen in its dendrogram. Trimmed K-means was used for subsequent characterization because it combined the best separation among the clusters without the influence of outliers.

**Table 1:** Calinski–Harabasz Index (CHI) for *K* = 5

| **Method** | **CHI** |
|---|---|
| K-means | 0.3186 |
| Trimmed K-means | 0.3308 |
| Single linkage | 0.0783 |
| Complete linkage | 0.2584 |
| Average linkage | 0.1755 |
| Ward linkage | 0.2742 |

The retained trajectories formed five recognizable mean temporal profiles under trimmed K-means (Fig. 4). Cluster 1 showed the highest maximal filopodia length. Cluster 2 is differentiated by the rate of protrusion and retraction. Cluster 3 has a fast extension rate to an early maximum and then retracted progressively. Cluster 4 followed a broad, approximately unimodal profile with a maximum near the first half of the lifetime and not alot of local undulation. Cluster 5 was more heterogeneous and showed two mean maxima, one near the middle and another near the end of normalized lifetime. It had the most local undulation. The colored means represent the predominant shapes rather than patterns followed identically by every member.

**Figure 4:**
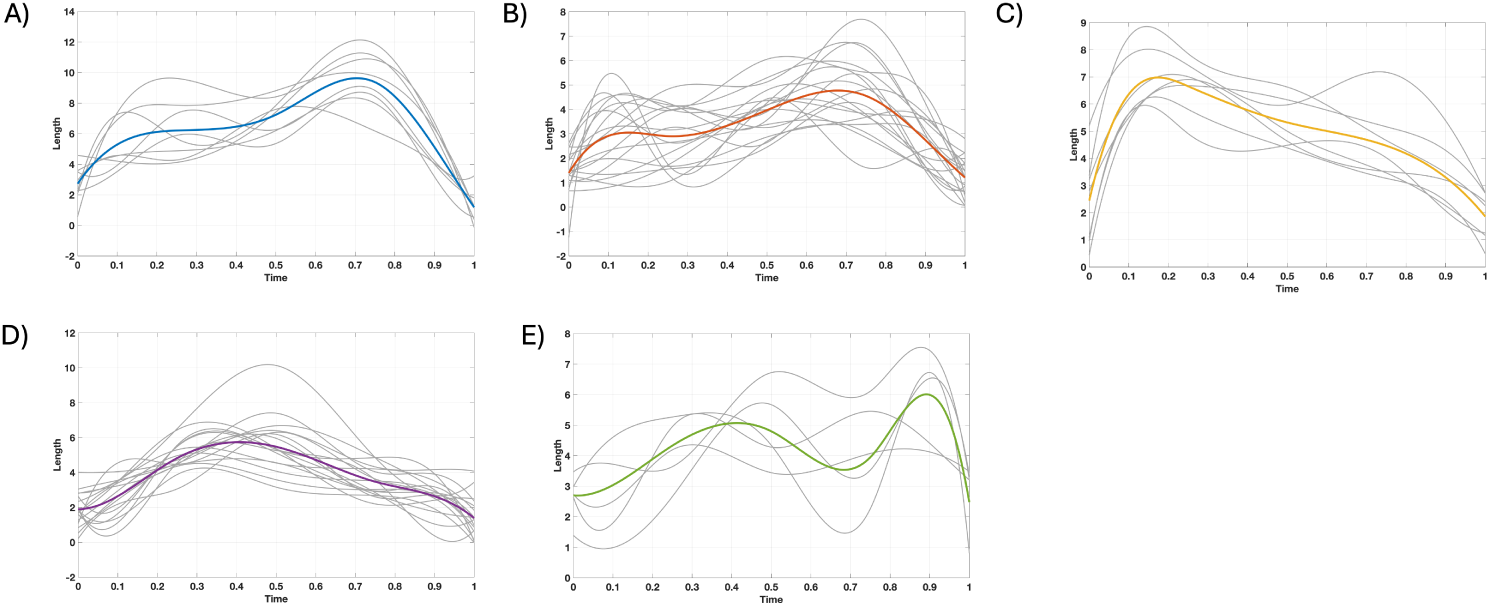
Filopodial length profiles in the five-cluster trimmed K-means solution. Gray curves show the fitted trajectories assigned to each cluster, and the colored curve shows the cluster mean. Panels correspond to Clusters 1–5 left to right. The clusters distinguish a late second-peak profile (Cluster 1), a lower-amplitude broad late peak (Cluster 2), an early-peak profile followed by sustained retraction (Cluster 3), a broad unimodal profile (Cluster 4), and high oscillatory profile (Cluster 5). Time is normalized within each filopodium.

Pairwise score projections clarify which FPCA modes drove the partition (Fig. 5). Cluster 1 was separated most clearly by its negative PC1 scores, consistent with a trajectory contrast captured by the dominant component. Cluster 5 occupied distinctly high PC4 scores and was therefore primarily distinguished by the localized variation represented by PC4. Clusters 2–4 overlapped in several two-dimensional projections, although Cluster 3 tended toward lower PC3 and PC4 scores and Cluster 4 tended toward lower PC2 scores. Thus, separation depended on the joint four-dimensional score space and cannot be inferred completely from any single two-component plot. The three trimmed trajectories fell outside or near the margins of the principal cluster regions in several projections, consistent with their large nearest-centroid distances.

**Figure 5:**
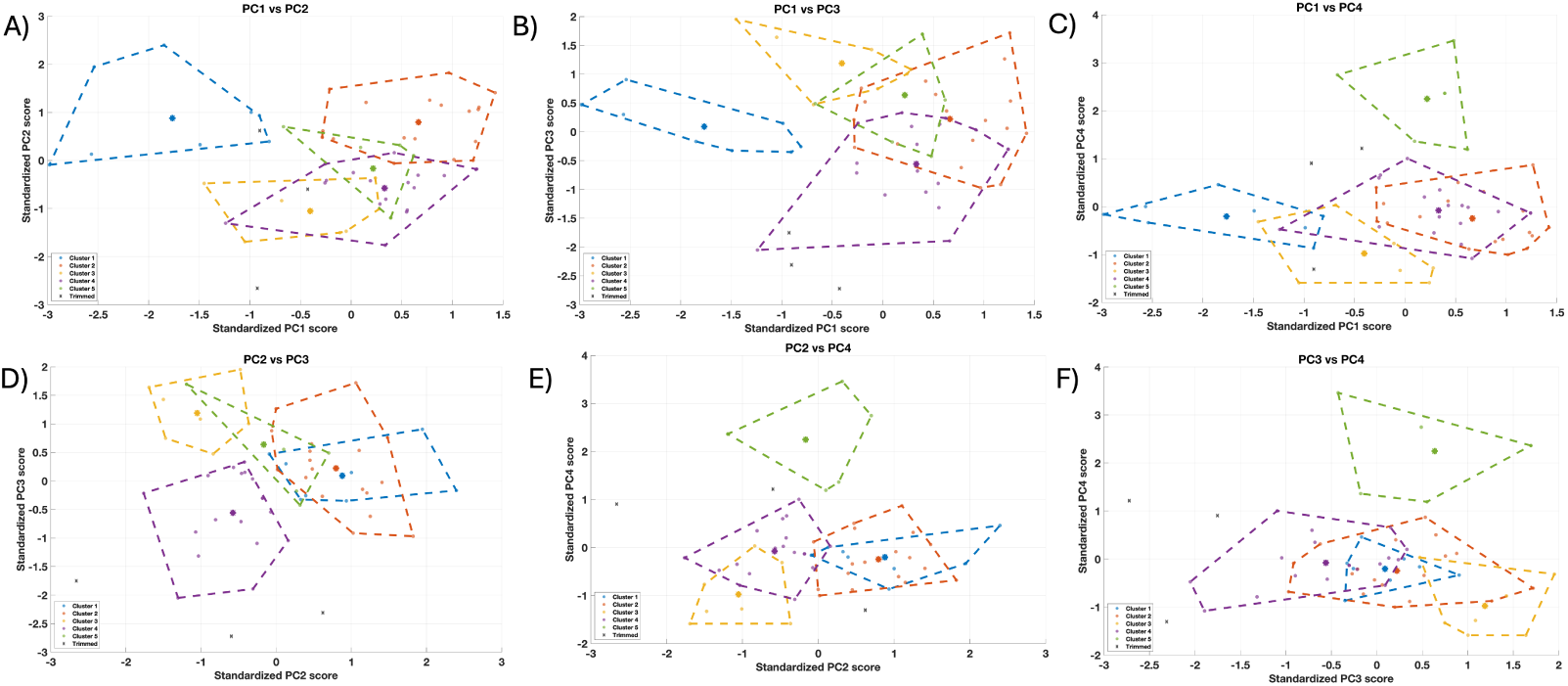
Pairwise projections of standardized FPCA scores for the trimmed K-means solution. Colored circles denote retained filopodia, dashed polygons outline the two-dimensional convex hull of each cluster, and asterisks mark projected cluster centroids. Black crosses identify the three trajectories excluded by 5% trimming. The six panels show all pairwise combinations of PC1–PC4. Cluster 1 is most clearly separated along PC1 and Cluster 5 along PC4, whereas Clusters 2–4 overlap in several projections, demonstrating that membership is determined in the complete four-dimensional score space. Convex-hull overlap in a projection does not imply overlap in all four dimensions.

**Figure 6:**
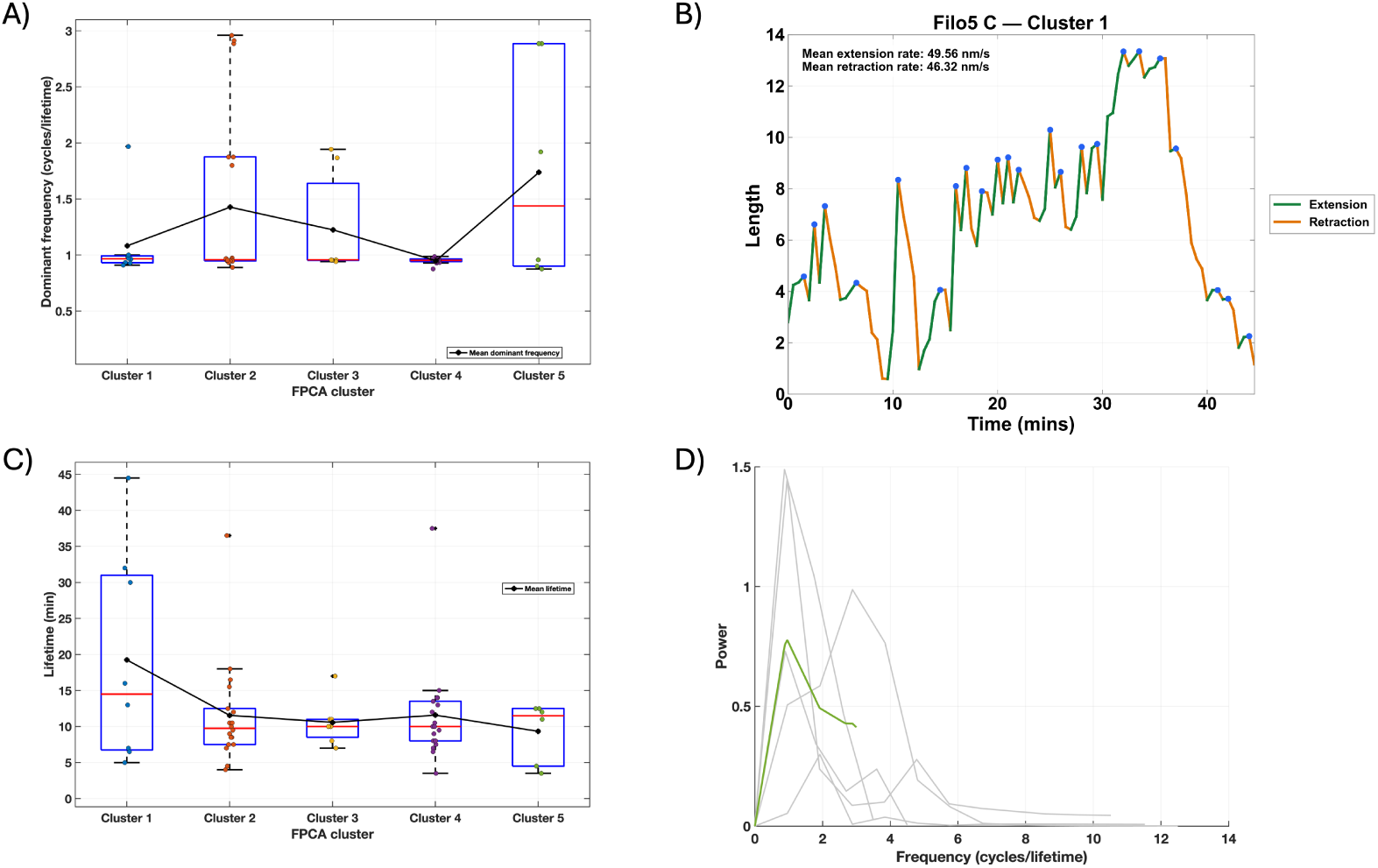
Dynamic and temporal characteristics of the filopodial clusters.(A) Distribution of dominant frequency across the five FPCA clusters,expressed as cycles per observed lifetime. Individual points represent filopodia, boxes show the interquartile range, red lines indicate medians,and connected black markers indicate cluster means.(B) Representative filopodial trajectory illustrating the calculation of mean extension and retraction rates. The green and orange lines indicates all extension and retraction event respectively over the course if the filopoduim’s lifetime, while the blue dot indicate the end of a extension event and the beginning of a retractione event.Rates were calculated using the original observation times. (C) Distribution of observed filopodia lifetime across clusters.Individual points represent each filopodia, boxes show the interquartile range,red lines indicate medians, and connected black markers indicate cluster means. (D) One-sided power spectra of the filopodia assigned Cluster 5. Gray curves represent individual filopodia, and the green curve represents the cluster-average spectrum Most spectral power occurs at low frequencies, although the individual spectra show substantial variability.

### 3.3. Characterization of filopodia clusters

Cluster characterization was performed on the original length and physical-time scales to recover biological differences that relative-time normalization intentionally removed. The five shape-defined clusters showed descriptive differences in size, persistence, rate, and fluctuation frequency (Table 2). Cluster 1 reached the greatest maximum length (9.78 ± 1.54 µm) and had the longest mean lifetime (19.25±14.54 min) and fastest mean extension rate (61.20 ± 34.22 nm s^-1^) and retraction rate (50.89 ± 22.85 nm s^-1^). Cluster 2 had the slowest mean extension rate (31.93 ± 13.66 nm s^−1^). Cluster 5 had the shortest mean lifetime (9.33 ± 4.17 min) and the highest mean dominant frequency (1.7368 ± 0.9716 cycles per lifetime).

**Table 2:**
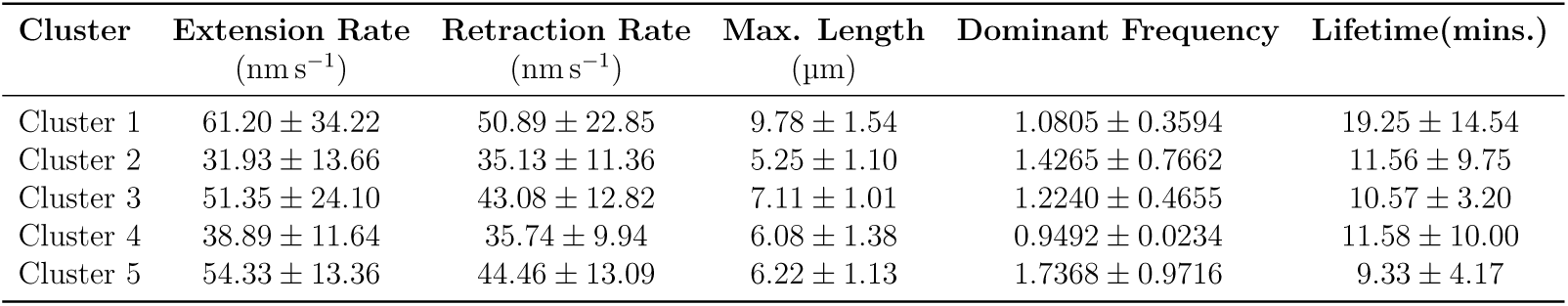
Physical and frequency-domain characteristics of the five trimmed K-means clusters. Values are mean ±standard deviation across retained filopodia assigned to each cluster. Extension and retraction rates were calculated from the original measurements and observation times on either side of the observed maximum; retraction is reported as a positive magnitude. Maximum length and lifetime were also calculated from the original records. Dominant frequency is expressed as cycles per observed lifetime. Undefined extension or retraction rates for trajectories peaking at the first or final observation, respectively, were omitted from the corresponding rate summary.

## Discussion

In this study, we applied functional data analysis and clustering techniques to characterize the temporal dynamics of filopodia using their complete length trajectories.Our results show that filopodia have unique and recurring patterns of retraction and extension that are not adequately explained by standard scalar measurements. 94.29% of the total trajectory variance was explained by the first four functional principal components, with overall length accounting for the largest percentage. This was followed by variations in the timing of extension and retraction, stabilizing behavior, and localized undulations.Subsequent application of trimmed K-means clustering identified five characteristic temporal profiles that differed in maximum length, lifetime, extension and retraction rates, and frequency-domain behavior. These findings are consistent with previous experimental studies demonstrating that filopodia are highly dynamic structures whose behavior involves multiple phases of growth, stabilization, and retraction[30].

One of the principal findings of this study is that overall filopodial length is the primary source of variation among the observed trajectories. The first functional principal component, which primarily captured differences in trajectory amplitude, accounted for 56.83% of the total variation. This finding is consistent with previous experimental studies that have emphasized filopodial length as an important measure of protrusive behavior [17]. The significance of length as a key characteristic of filopodial behaviour is confirmed by our results. They further show that the diversity of filopodial dynamics cannot be fully captured by length-related variation alone. Together, the other three principal components explained an additional 37.47% of the variation, even though the first principal component explained the most amount. These elements explained variations in localised undulation, duration of stationary phase, and the timing of extension and retraction. This distinction is especially crucial, because filopodia with similar maximum lengths may display very different varying temporal trajectories. For example, one filopodium may quickly reach its maximum length and retract right away, whereas another may gradually achieve the same length and stay relatively extended until retracting. Therefore, a description based only on maximum length would not be able to differentiate between these behaviours. Hence, our analysis broadens earlier quantitative research by revealing that the comprehensive temporal trajectory can facilitate the identification of diverse dynamic profiles without initially simplifying each filopodium to a limited range of scalar indicators.

## Conclusion

In this work, we used the full length-versus-time trajectories of filopodia to examine their temporal dynamics utilizing functional data analysis and clustering techniques. Through the use of trimmed K-means clustering, cubic B-spline representations, and functional principal component analysis, we were able to identify five recurrent dynamic characteristics in 60 filopodia that were observed. 94.29% of the entire trajectory variance was explained by the first four functional main components, with overall length making up the biggest percentage. This was followed by variations in the timing of extension and retraction, stabilizing behavior, and localized fluctuations. Our results show that while maximal length is the primary cause of variation in filopodial dynamics, variation of their temporal behavior is not entirely captured by it. Maximum length, observed lifetime, time-weighted extension and retraction rates, and frequency-domain features vary among the five detected clusters. These variations show how crucial it is to take into account filopodia’s entire temporal evolution rather than depending solely on scalar measurements. The findings are in line with earlier experimental research showing that filopodia experience intricate patterns of retraction, stability, and extension. Our method, however, goes beyond these findings by offering a methodical, data-driven framework for locating recurrent temporal profiles without predetermining the anticipated dynamic patterns. The identification of distinct dynamic classes also provides a foundation for investigating the relationship between filopodial behavior and long-range Notch signaling. In particular, differences in spatial reach, stabilization, and repeated extension–retraction events may influence opportunities for receptor–ligand interactions and mechanical force transmission. Establishing these relationships will require future studies combining detailed trajectory measurements with mathematical models, measurement of mechanical forces, and Notch activation. Overall, this study demonstrates the value of functional data clustering for characterizing the heterogeneity of filopodial dynamics and provides a quantitative framework for future investigations linking temporal behavior to cellular signaling and biological function.

## Author Contributions

NE wrote the original draft of the manuscript, performed the functional data analysis, including FPCA, data preprocessing, clustering and statistical characterization of filopodia dynamics. SHDSDS, HC, GS tracked filopodia trajectories and performed preliminary length analysis. G.H designed and supervised all experiments, jointly conceptualized the study with EAA, and reviewed manuscript drafts. EAA conceptualized study, designed methods, supervised the interpretation of results and reviewed initial drafts of the manuscript.

## Acknowledgments

This project is supported by the joint NSF/NIGMS mathematical biology program. Award number R01GM152810. Work in the Hunter lab is supported by R35GM150782.

